# Fast 2-photon stimulation using holographic patterns

**DOI:** 10.1101/2023.01.12.523269

**Authors:** Maia Brunstein, Jules Lubetzki, Cédric Moutoussamy, Wei Li, Jérémie Barral

## Abstract

Two decades after its introduction, optogenetics – a biological technique to control the activity of neurons or other cell types with light – remains a cutting edge and promising tool to study biological processes. Its increasing usage in research varies widely from causally exploring biological mechanisms and neural computations, to neurostimulation and sensory restauration. To stimulate neurons in the brain, a variety of approaches have been developed to generate precise spatiotemporal light patterns. Yet certain constrains still exists in the current optical techniques to activate a neuronal population with both cellular resolution and millisecond precision. Here, we describe an experimental setup allowing to stimulate a few tens of neurons in a plane at sub-millisecond rates using 2-photon activation. A liquid crystal on silicon spatial light modulator (LCoS-SLM) was used to generate spatial patterns in 2 dimensions. The image of the patterns was formed on the plane of a digital micromirror device (DMD) that was used as a fast temporal modulator of each region of interest. Using fluorescent microscopy and patch-clamp recording of neurons in culture expressing the light-gated ion channels, we characterized the temporal and spatial resolution of the microscope. We described the advantages of combining the LCoS-SLM with the DMD to maximize the temporal precision, modulate the illumination amplitude, and reduce background activation. Finally, we showed that this approach can be extended to patterns in 3 dimensions. We concluded that the methodology is well suited to address important questions about the role of temporal information in neuronal coding.

## Introduction

A fundamental question in neuroscience is to understand how patterns of neuronal activity can represent useful sensory or motor information to drive relevant animal behavior. To build a causal link between brain activity and behavior, it is necessary to control the neuronal dynamics and observe how animal comportment is affected (Panzeri, Harvey et al. 2017). Addressing this challenge requires tools to manipulate a population of neurons in a very controlled manner, both spatially and temporally. This can now be achieved by optogenetics, a genetic manipulation that drives the expression a light-modulated actuator in a defined cell type in order to modify cellular properties during illumination. These actuators interact with diverse biochemical pathways (transcription, second messengers, or other signaling pathways) or can be ion pumps or channels (Emiliani, Entcheva et al. 2022). The latter category is particularly interesting for neuroscience applications because neurons expressing the light gated ion channel channelrhodopsin or alternative variants can be efficiently activated by brief light pulses (Yizhar, Fenno et al. 2011). Combined with diverse optical methods to produce precise light patterns, it allows to specifically activate a defined ensemble of neurons (Ronzitti, Ventalon et al. 2017, Chen, Papagiakoumou et al. 2018). Because neurons are embedded in the scattering brain tissue, 2-photon (2P) activation is often necessary to reach cells in deep structures with single-cell resolution. Two classes of strategies have been developed to stimulate neurons using 2P activation. A first possibility is to scan a femtosecond infrared laser beam onto target cells using galvanometric mirror (Rickgauer and Tank 2009) or optoacoustic deflectors (Ricci, Marchetti et al. 2022). However, this approach is too slow to activate a large population of neurons with good temporal precision, especially in 3 dimensions.

The optimal choice for stimulating simultaneously several cell targets is to benefit from non-scanning methods (Papagiakoumou, Anselmi et al. 2010, Papagiakoumou, Ronzitti et al. 2018), such as holography which enables to generate arbitrary patterns in 3 dimensions. However, using this technique alone to update temporal patterns rapidly is limited by two major bottlenecks. First, the refresh rate of the fastest Liquid Crystal on Silicon Spatial Light Modulator (LCoS-SLM) on the market is effectively limited to about 300-400 Hz because liquid crystals have a rise time of ∼2 ms when used with infrared wavelengths. Therefore, better temporal precision cannot be achievable in this configuration. The second constraint concerns the number of phase patterns to generate: if for example we need to stimulate several targets with random temporal patterns, one can reasonably assume that each pattern of the sequence will be different. Considering a 1 s stimulation sequence with a 500 Hz refresh rate of the LCoS-SLM, one will need to build 500 phase patterns. Although this is feasible to pre-build the patterns and store them either on a computer or on the onboard memory of the device, it precludes any real-time control of neuronal activity because of the necessary computational time to calculate these patterns with the Gerchberg-Saxton iterative algorithm (Gerchberg and Saxton 1972).

Considering 1P stimulation, one simple approach is to use a spatial light modulator in a plane conjugated with the sample plane and modulate the amplitude of the light (Ronzitti, Ventalon et al. 2017). An example of spatial modulator is a Digital Micromirror Devices (DMD) which can achieve binary frame rates up to 10 kHz. This device has been successfully employed to activate an arbitrary number of targets with sub-millisecond precision (Barral and Reyes 2016, Barral and Reyes 2017). However, the intensity of each pixel is, in this case, the intensity of the light source divided by the total number of pixels. As a simplified example and taking a very conservative approach, if a 40 W laser is able to bring 10 W of light to the sample, we cannot expect more than 10 μW of light per pixel with a DMD of 1000×1000 pixels. With a pixel size on the sample of 0.5 μm (considering a 20X magnification and a 10 μm pixel size on the DMD), it represents 40 μW·μm^-2^. This is clearly enough for 1P stimulation where the saturating activation of channelrhodospin ChR2 is usually around 0.01 μW·μm^-2^, but this only allows to activate a small fraction of opsin molecules with 2P stimulation (Andrasfalvy, Zemelman et al. 2010, Pegard, Mardinly et al. 2017, Chen, Ronzitti et al. 2019) (see also Fig. 4e below). To reliably activate the opsin and reach a light intensity of 100 μW·μm^-2^ necessary for scanless 2P activation with this scenario, we would need to increase the laser power to at least 120 W. This approach is thus both inefficient because of the loss of most of the light power and unfeasible because such a laser is not presently available.

To overcome these limitations, we used a LCoS-SLM to generate a single spatial pattern in 2 or 3 dimensions containing multiple regions of interest and projected this pattern onto a DMD that was employed as a fast temporal shutter for each region of interest. In this way, the total laser power was efficiently dedicated to activating only potential targets without wasting laser power on unused DMD pixels. This approach had been applied to imaging application with the development of an encoded multisite 2P microscope (Ducros, Goulam Houssen et al. 2013) or to multi-site photolysis of caged neurotransmitters (Go, To et al. 2013). The main advantages of our optical developments are (i) to push the temporal resolution to the limitations of the opsin time constant, (ii) to reduce the complexity of the phase pattern calculation since only 1 phase pattern needs to be built, (iii) to enable real-time control of neuronal activity. We further demonstrate that the set-up allows to generate patterns with increased contrast and enables stimulation in 3 dimensions.

## Methods

### Microscope apparatus

To achieve a precise control of the light pattern both temporally and spatially, we developed a homemade high-speed photo-stimulation system coupled to a 2-photon microscope (Fig 1). The architecture of the 2-photon microscope was inspired by an open-source design previously published (Rosenegger, Tran et al. 2014). The excitation beam (920nm, Opera HP, Coherent) was first sent through a λ/2 wave retarder (AHWP10M-980, Thorlabs) mounted on a motorized rotation stepper motor (K10CR1/M, Thorlabs) and a polarizer cube (GL10-B, Thorlabs) for control of the laser power. The beam was expanded through a telescope (f_1_ = 40 mm, AC254-040-B, Thorlabs; f_2_ = 60 mm, AC254-060-B, Thorlabs) and sent to a pair of scanning galvanometer mirrors (GVSM002/M, Thorlabs). After the scanner, the beam impinged the scan lens (LSM04-BB, Thorlabs) followed by the tube lens (f = 200 mm; AC508-200-B, Thorlabs) to be finally focalized by a 20X objective (XLUMPLFLN20XW, Olympus). The emission of the sample was collected with the same objective and sent via a dichroic filter (DMLP735B, Thorlabs) through an emission tube lens (f = 150 mm, AC254-150-A, Thorlabs) to the detection path. The detection unit was composed of a dichroic filter (DMLP567R, Thorlabs) to separate the green and the red channels, each of them were de-scanned via a collector lens (f = 30 mm, LA1805-A, Thorlabs), filtered (FF01-520/70-25 and FF01-650/150-25 for the green and the red channels, respectively, Semrock) and detected by a photomultiplier tube (GaAsP PMT, PMT2101/M, Thorlabs, and Multialkali PMT, PMT1001/M, Thorlabs). In order to find the region of interest on the focal plane, we added an epi-fluorescence system, composed by a 470 nm LED, a 469/35 excitation filter, a 500LP dichroic, and a 525/40 emission filter (MDF-GFP, Thorlabs). Because our laser for 2P microscopy had a low repetition rate (2 MHz), we synchronized the acquisition of the fluorescent signal with the laser pulses to correct the fluorescent signal by the actual number of laser pulses within each pixel. To do so, we built a custom user interface programmed with LabVIEW (National Instrument) that controlled the whole microscope apparatus described here.

**Fig. 1.**
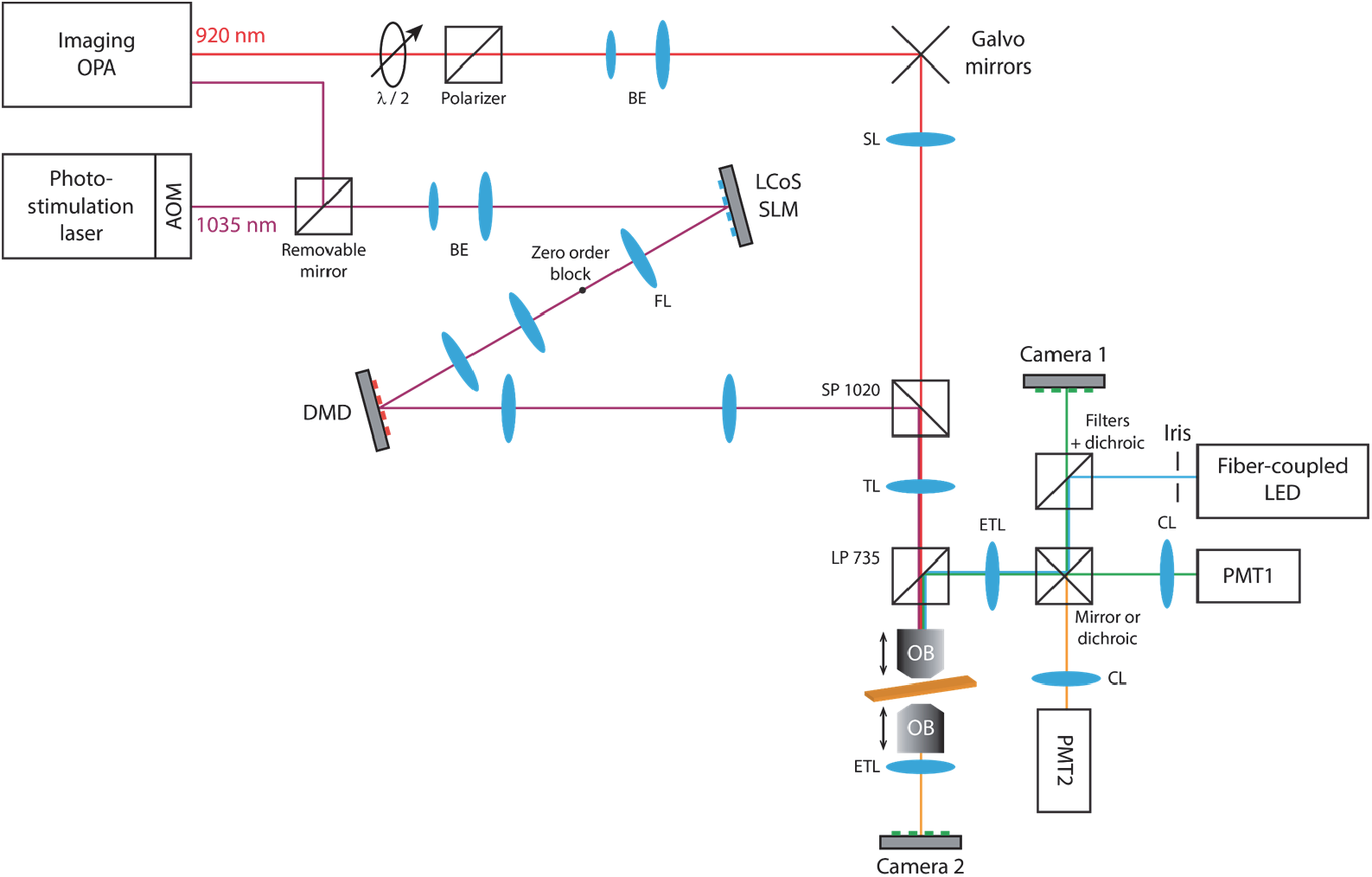
Optical light paths of the 2-photon microscope and 2-photon stimulation device. The 920 nm laser was scanned on the sample plane trough the objective by the galvanometer mirrors. The fluorescence emission was collected by the same objective and sent to the PMT. The phase of the 1035 nm laser was modulated by the LCoS-SLM to form an arbitrary pattern on the DMD surface. This last element was used as a fast temporal shutter to excite different regions of interest on the sample plane with high temporal resolution. A second objective was placed under the sample to image patterns in 3D. Abbreviations: AOM: acousto-optic modulator; BE: beam expander; CL: collector lens; DMD: digital micromirror device; ETL: emission tube lens; FL: Fourier lens; LCoS-SLM: liquid crystal on silicon spatial light modulator; LP: long-pass dichroic mirror; OB: objective; OPA: optical parametric amplifier; PMT: photomultiplier tube; SL: scan lens; SP: short-pass dichroic mirror; TL: tube lens.

In the photostimulation path, the emission of the laser (1035nm, Monaco, Coherent) was expended through a telescope (f_1_ = 30 mm, AC254-030-B, Thorlabs; f_2_ = 250 mm, AC508-250-B, Thorlabs) to fill the sensitive area of the LCoS-SLM (E-series 1920×1152 pixels, Meadowlarks optics). The LCoS-SLM allowed modifying the phase of the incident beam at the back focal plane of the objective in order to create an arbitrary intensity pattern on the image plane (i.e. several diffraction-limited spots, large patterns or arbitrary shapes in 3D). The beam reflected from the LCoS-SLM formed the first intensity pattern in the focal plane of the Fourier hologram lens (f = 150 mm, AC508-150-B, Thorlabs). A blocking mask in this plane was used to obstruct the ‘zero order’ of the LCoS-SLM (non-modulated light). To be able to modify the illumination of individual targets at high speed, we relayed this intensity pattern to a DMD (DLP650LNIR, 1280×800 pixels, Texas Instruments) via a 1:1 telescope (f = 250 mm, AC508-250-B, Thorlabs). The DMD was controlled by a third-party controller (V-650L, Vialux) to achieve full-frame temporal modulation above 10 kHz. Finally, the phase-modulated beam was relayed using a 1:1 telescope (f = 250 mm, AC508-250-B, Thorlabs) and combined to the two-photon excitation beam with a 1020 nm short-pass dichroic mirror (DMSP1020B, Thorlabs) before going through the tube lens and the objective and being projected on the sample plane.

To characterize 2D patterns, we used a camera located at Camera 1 position. For the analysis of contrast ratio, we visualized the reflected fluorescence emitted from a thin layer of fluorescent ink enclosed between a slide and a coverslip. A CMOS camera (acA1920-40um, Basler) was placed after the emission tube lens and a short-pass filter (FESH0750, Thorlabs). To characterize the spatiotemporal properties of the optical setup, we visualized 2D patterns generated onto a fluorescent microscope slide (Chroma) placed at the sample plane and imaged through the same fluorescent light path by a rapid camera (ORCA-Flash4.0, Hamamatsu).

To reconstruct 3D patterns, a visualization system composed of an additional 20X objective (UMPLFLN20XW, Olympus), a short-pass filter (FESH0750, Thorlabs) a tube lens (f = 150 mm, AC254-150-A, Thorlabs) and a CMOS camera (acA1920-40um, Basler) was positioned under the sample (Camera 2 position). The thin layer sample was positioned at the focal plane of the lower objective and both the sample and the lower visualization unit were mounted onto a motorized translation stages (PLS-X, Thorlabs). This configuration was used to visualize the 3D illumination patterns as well as the intensity profiles of the stimulation spots in X-Y-Z.

### Calibration and image analysis

The system was calibrated before each use to match the image from the camera (Camera 1) with both the LCoS-SLM and DMD projection units and the image from the 2P microscope. For simplicity, the image from the camera was considered as the reference. To calibrate the LCoS-SLM, a phase mask was sent from the LCoS-SLM to create a grid of 4×4 spots while the DMD was in an all-ON configuration. Spots generated by the LCoS-SLM were detected on the camera and a geometric matching algorithm (Vision Development Module, LabView, National Instruments) was used to calibrate for perspective or nonlinear distortion errors. With our settings, the LCoS-SLM pixel size was 0.43 μm x0.72 μm at the sample plane. To calibrate the DMD, the LCoS-SLM sent an uniform illumination pattern to the DMD which was configured to create a grid of 4×4 spots. With our settings, the DMD pixel size was 0.50 μm x0.53 μm at the sample plane. It should also be noted that the image of the LCoS-SLM at the DMD plane (which was a square of 16.8 mm side) was larger than the DMD active area (13.8 mm x 8.6 mm), such that the field of excitation was only defined by the locations accessible with the DMD and took a value of 640 μm x 430 μm at the sample plane. Finally, the 2P imaging path was calibrated by sending spots sequentially at different locations using fixed angle deflections with the galvanometer mirrors.

We also used the calibration grid generated by the LCoS-SLM to adjust the light intensity of the different spots. Because light is diffracted differently according to the target location, we first fitted the intensity profile of the spots in the (*x,y*) plane by a 2-dimensional Gaussian function and compensated the target spot intensity to achieve a nearly homogenous field of excitation. This simple method allowed us to decrease the standard deviation of the 2P absorption from 42% down to 18% (Fig. 2) comparable to published data (Hernandez, Papagiakoumou et al. 2016, Pegard, Mardinly et al. 2017).

**Fig. 2.**
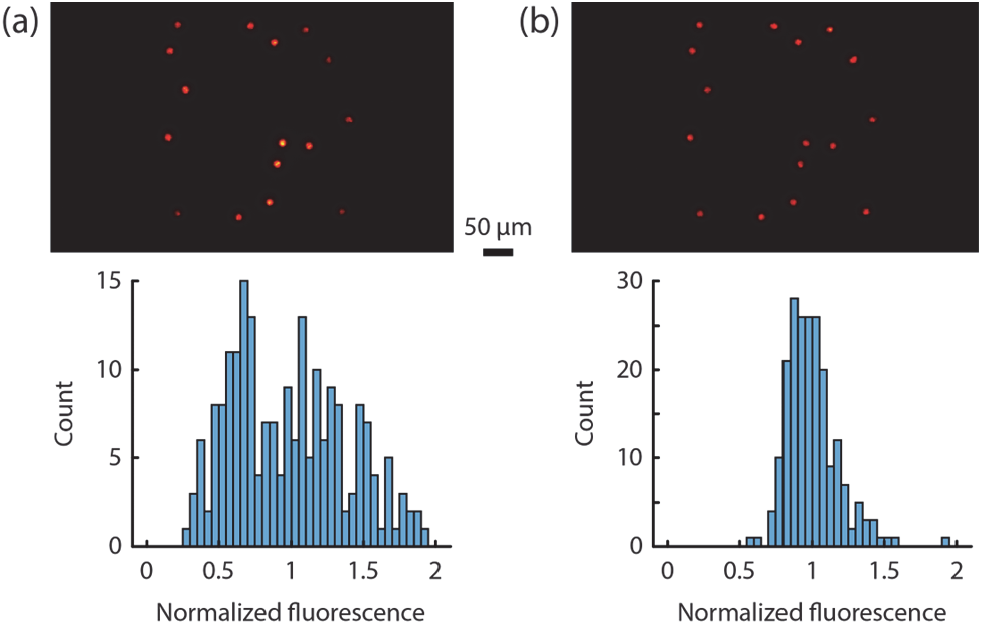
Diffraction efficiency characterization and correction. (a) Example of an optical pattern containing 16 disks of 10 μm diameter randomly located in the field of excitation without any correction for intensity (top). Histogram of the normalized integrated intensity of the spots for different patterns (data were collected from 10 different patterns with 16-32 ROIs randomly positioned in the field of excitation). (b) Same as in (a) after intensity compensation.

Image analysis and quantification were performed with the Image Processing toolbox (Matlab). Three-dimensional visualization was performed with Vaa3d software (Peng, Ruan et al. 2010) or with Matlab.

### Neuronal cultures and electrophysiology

Dissociated cortical neurons from postnatal (P0-P1) mice were prepared as described previously (Hilgenberg and Smith 2007, Barral and Reyes 2016, Barral and Reyes 2017) and in accordance with French and European regulations for the care and protection of laboratory animals (EC Directive 2010/63, French Law 2013–118). Briefly, the mouse cortex of FVB wild type mice (Janvier Laboratory) was dissected in cold CMF-HBSS (Ca^2+^ and Mg^2+^ free Hank’s balanced salt solution containing 1 mM pyruvate,15 mM HEPES, 10 mM NaHCO_3_). The tissue was dissociated in papain (15 U/mL, Roche) containing 1 mM L-cystein, 5 mM 2-amino-5-phosphonopentanoic acid and 100 U/ml DNase (DN25, Sigma) for 25 min. After enzymatic inactivation in CMF-HBSS containing 100 mg/mL BSA (A9418, Sigma) and 40 mg/mL trypsin inhibitor (T9253, Sigma), pieces were mechanically dissociated with a pipette and plated on a glass coverslip. Before plating, glass was cleaned in 3 N HCl for 48 h and immersed in sterile aqueous solution of 0.1 mg/mL poly-L-lysine (P4707, Sigma) for 12 h. Neurons were grown in Neurobasal medium (supplemented with B27, Glutamax and penicillin/streptomycin cocktail, Gibco) in a humidified incubator at 37 °C, 5% CO_2_. One third of the culture medium was exchanged every 3 days.

Expression of channelrhodopsin (ChR2) was achieved by viral infection with AAV2-hSyn-hChR2(H134R)-EYFP of cortical neurons. The virus was produced at 3×10^12^ cfu/mL by the University of Pennsylvania Vector Core using plasmid provided by Karl Deisseroth (Stanford University). At 3 days *in vitro* (DIV), the culture was infected with 1 μL of virus. Experiments were performed at 14-21 DIV, when neuronal characteristics and network connectivity were mature and expression of ChR2 was sufficient to enable reliable photostimulation.

Recordings were performed at room temperature in artificial cerebrospinal fluid (aCSF). The aCSF solution contained (in mM): 125 NaCl, 15 NaHCO_3_, 25 D-glucose, 10 HEPES, 2.5 KCl, 2 CaCl_2_, 1.25 NaH_2_PO_4_ and 1 MgCl_2_. Electrodes, pulled from borosilicate pipettes (1.5 OD) on a Flaming/Brown micropipette puller (Sutter Instruments), had resistances in the range of 6-10 MΩ when filled with internal solution containing (in mM): 130 K-gluconate, 10 HEPES, 10 phosphocreatine, 5 KCl, 1 MgCl_2_, 4 ATP-Mg and 0.3mM GTP.

Cells were visualized through the water-immersion objective and fluorescence microscopy was used to confirm ChR2 expression. Whole-cell current-clamp recordings were made using Intan CLAMP amplifier (Intan Technologies). The signal was filtered at 10 kHz and digitized at 50 kHz.

## Results

### Microscope design and pattern quality

The 2P stimulation path of the microscope was based on a fixed-wavelength femtosecond laser at 1035 nm capable of delivering a power of 40 W (Fig. 1). The laser beam was expanded onto the LCoS-SLM and the zero-order diffraction was physically eliminated. The image was then projected onto the DMD and further relayed to the sample plane. Several calibration steps were necessary to find the corresponding locations of pixels on the LCoS-SLM, DMD, camera, and 2P scanning image (see Methods). Once the system was calibrated, the user defined Regions of Interest (ROIs) on the sample that was imaged either by the camera or by the 2P microscope. Typically, ROIs were disks of 10 μm diameter at the sample plane which corresponds to the average size of a neuron in neuroscience applications. The corresponding amplitude masks in the LCoS-SLM and DMD coordinate systems were then computed. A unique phase mask containing all the ROIs was generated using the Gerchberg-Saxton iterative algorithm (Gerchberg and Saxton 1972) and projected by the LCoS-SLM. A collection of corresponding DMD masks were also designed. Each mask defined whether each ROI was relayed or not to the sample. The DMD was thus used as a switchable mirror for each region of interest. Because theses masks were simply 1-bit images, they could be pre-built or generated and uploaded rapidly for real-time applications.

Some concerns could be raised whether using only a fraction of the DMD mirrors could degrade the quality of patterns generated by the LCoS-SLM. To characterize the spatial performances of our photostimulation light path, we visualized with a camera the reflected fluorescence emitted from a thin layer of fluorescent ink enclosed between a slide and a coverslip. We measured the intensity profile in 3 dimensions of individual disks randomly distributed in the field of excitation when all mirrors of the DMD where in ON position (Fig. 3a) or when only the mirrors corresponding to the ROI were reflecting the optical pattern (Fig. 3b), while keeping the phase mask identical in both conditions. Both in the sample plane (Fig. 3c) or in a plane containing the optical axis (Fig. 3d), the different configurations provided similar profiles although having a mask on the DMD seemed to sharpen slightly the fluorescent pattern. Along the axial direction, the full width at half maximum (FWHM) decreased on average from 21.0 ± 0.3 to 20.0 ± 0.3 μm for a 10 μm disk (mean ± SE, n = 10, paired *t*-test with *p*-value < 10^−4^). A thickness of about 20 μm is very comparable to published data (Dal Maschio, Difato et al. 2010, Pegard, Mardinly et al. 2017, Shemesh, Tanese et al. 2017) but could notably be reduced with the introduction of temporal focusing (Papagiakoumou, de Sars et al. 2008, Andrasfalvy, Zemelman et al. 2010, Papagiakoumou, Anselmi et al. 2010, Pegard, Mardinly et al. 2017, Papagiakoumou, Ronzitti et al. 2020). In particular, temporal focusing would permit to avoid the decrease of spatial confinement as the size of the ROI is increased (Fig. 3e). We did not use this technique in the present study, but it would be adaptable to our optical design and would help to increase axial confinement if needed.

**Fig. 3.**
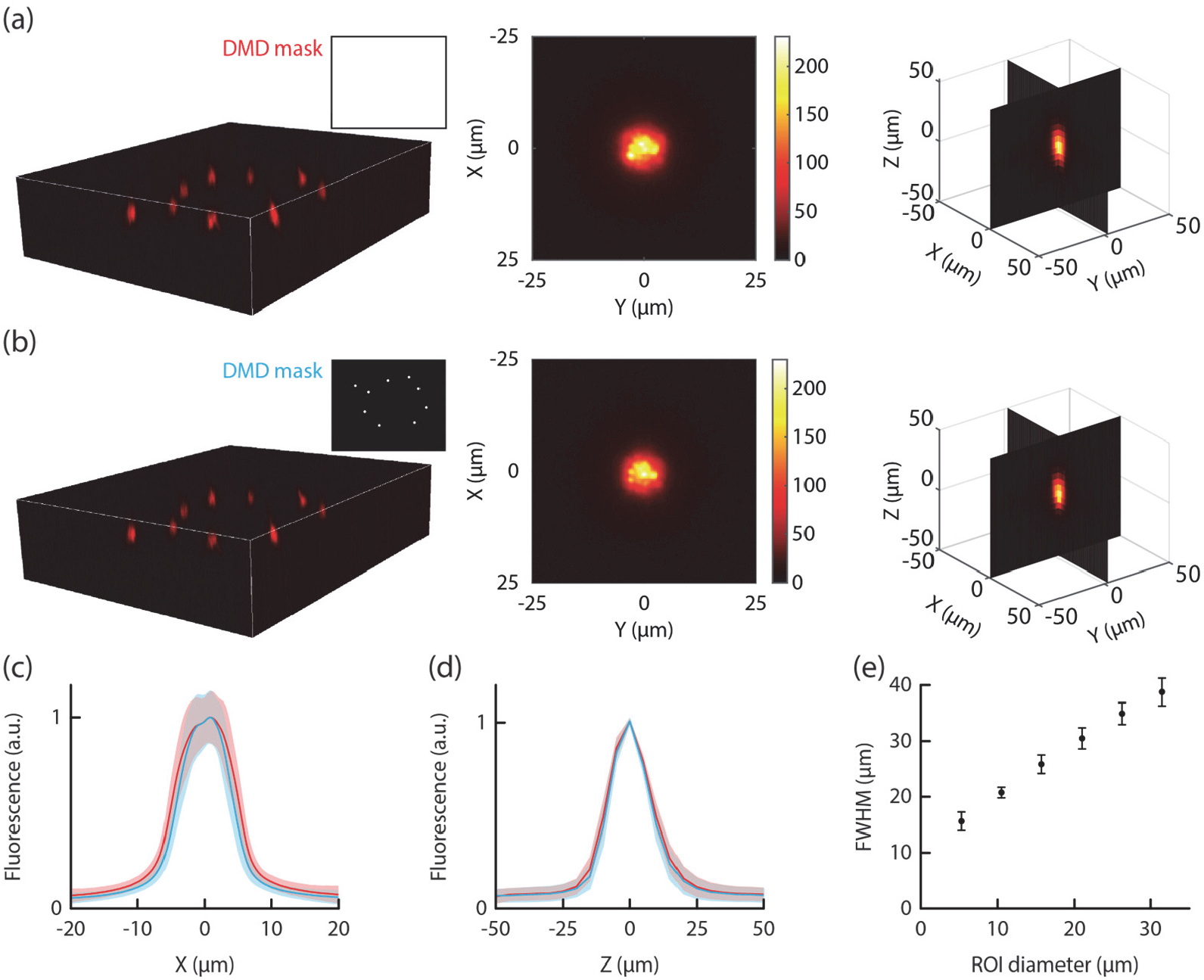
Profile of an optical pattern (10 μm diameter disk) when all mirrors of the DMD were in the ON position (a) or when only the mirrors corresponding to the ROIs were ON (b). 3-dimensional reconstruction of a stimulation volume (left). Intensity profiles of one stimulation target in the XY plane (middle) and in the planes along the optical axis (right). (c) Average (n = 75 ROIs) intensity profile along the X axis when all the mirrors of the DMD were in the ON position (red) or when only the mirrors corresponding to the ROIs were ON (blue). The shaded area represents ± 1 STD. Data were collected from patterns containing 5-20 ROIs simultaneously displayed in the same plane as in the examples provided in panels (a) and (b) and normalized to their maximal intensity. (d) Same as in (c) for the intensity profile along the Z axis (n = 9 ROIs). (e) Measured axial confinement (full width at half maximum, FWHM) of the 2P fluorescence intensity distribution produced by circular holographic spots of increasing size when 10 spots were simultaneously displayed (mean ± STD).

Altogether, these intensities profiles are comparable to those found in the literature using only a LCoS-SLM (Dal Maschio, Difato et al. 2010, Hernandez, Papagiakoumou et al. 2016, Papagiakoumou, Ronzitti et al. 2018) thereby suggesting that the introduction of a grid of micromirrors did not degrade significantly the quality of patterns generated by the LCoS-SLM.

### Improvement of contrast ratio

Comparing images acquired with a fully reflective DMD or when the DMD was only reflecting the corresponding ROIs shows that there is slightly less background light for equal target intensities (Fig. 3c). This can also be appreciated by subtracting the corresponding images (Fig. 4a): there is in proportion more light in the target than outside when the DMD is used as a spatial modulator. To fully characterize the amount of light projected to the desired area, we quantified the contrast ratio defined as the average pixel intensity within a ROI divided by the average background intensity. For this, we generated random patterns with 1-128 ROIs of 10 μm diameter each. When the pattern contained only a single ROI, the background intensity was low resulting in a high contrast ratio of ∼4900:1 (Fig. 4b) that was actually increased to ∼6200:1 when the DMD allowed to reflect only the corresponding ROI (Fig. 4c), resulting in an improvement by 25% (Fig. 4d). We then systematically increased the number of ROIs while adjusting the laser intensity (Fig. 4e) to maintain target’s intensities constant (Fig. 4b-c). Background intensity increased almost linearly, mirroring the increase of laser intensity that was necessary to keep the ROI’s intensity constant (Fig. 4e). This resulted in an inverse relationship between contrast ration and ROIs number (Fig. 4b-c, right). However, the contrast ratio with the DMD as a spatial modulator was consistently higher than the one obtained when the DMD was fully reflective, leading to an average improvement of contrast ratio by 21.8% (Fig. 4d).

**Fig. 4.**
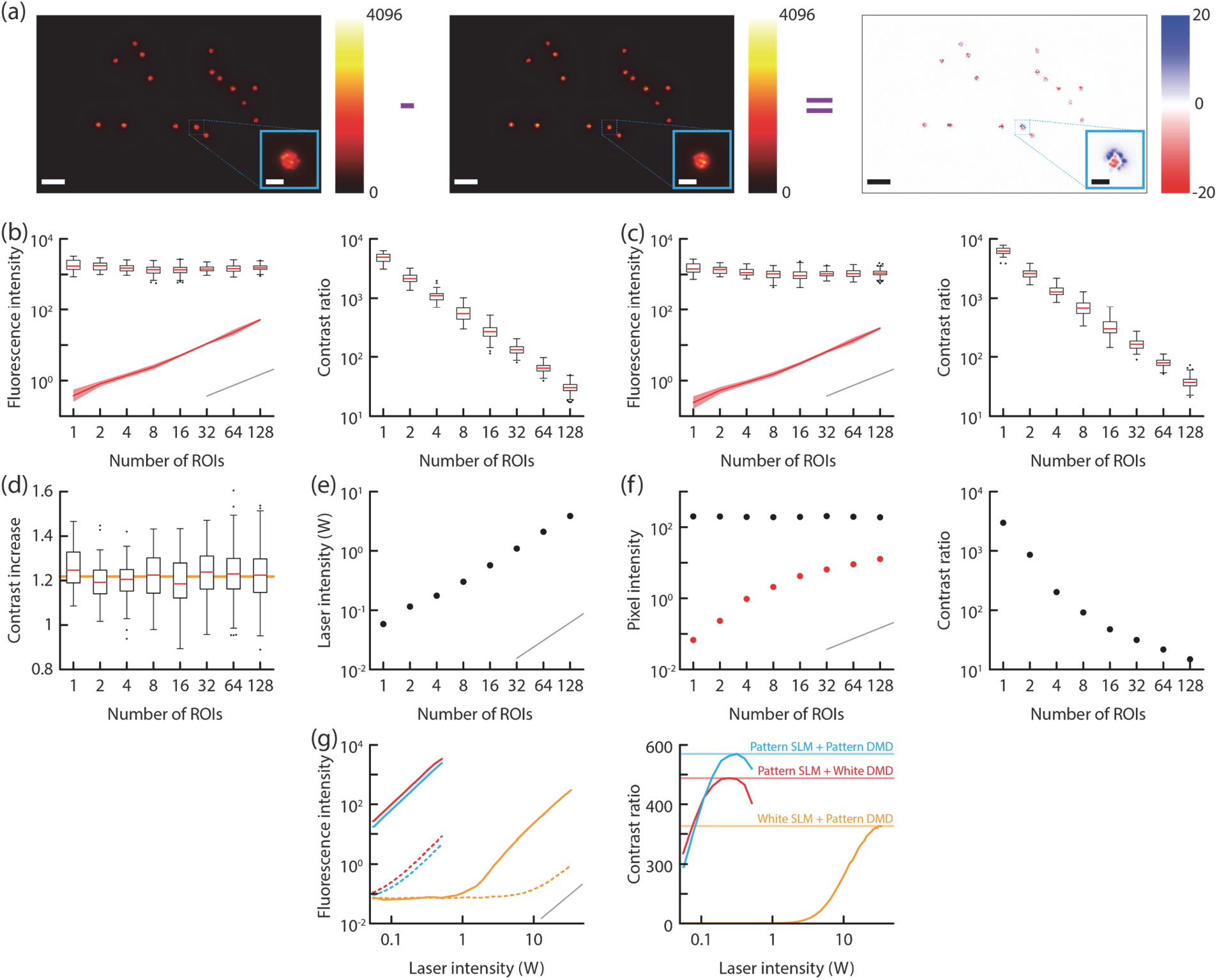
Contrast ratio of holographic patterns. (a) Example of an optical pattern containing 16 disks of 10 μm diameter randomly located in the field of excitation when all mirrors of the DMD were in the ON position (left) or when only the mirrors corresponding to the ROIs were ON (middle). Images were normalized to their mean intensity before being subtracted (right). Note that the inside of the targets shows relatively higher intensity (red) and that the border shows lower intensity (blue) when the DMD was turning OFF outside pixels. Scalebar 50 μm and 10 μm for the full image and the inset zoom, respectively. (b) left, Fluorescent intensity of the regions of interest (boxplot) and of the background (red shaded area representing the mean ± STD) as a function of ROIs’ number when all mirrors of the DMD were in the ON position. right, Corresponding contrast ratio. (c) Same as in (b) when only the mirrors corresponding to the ROIs were ON. (d) Increase in contrast ratio observed when only the mirrors corresponding to the ROIs were ON compared to a situation where the DMD was reflecting all the light. For each number of ROIs, the increase of contrast ratio is significantly higher than 1 (t-test statistic for each number of ROIs: P<10^−10^, n = 30-640, depending on ROIs number). The orange line represent the average increase. (e) Laser intensity used to match the mean ROI intensity when their number was increased. (f) left, Expected pixel intensity in the ROI (black) or in the background (red) computed from the holographic phase patterns. right, Corresponding contrast ratio. (g) left, Fluorescent intensity of the regions of interest (solid lines) and of the background (dashed lines) for a pattern containing 10 ROIs of 10 μm diameter as a function of laser intensity in 3 different conditions: when only the SLM was used to produce a pattern and the DMD was flat (red), when the SLM produced a pattern and the DMD relayed the same pattern (blue), or when the SLM was flat and only the DMD was used to produce the pattern (orange). right, Corresponding contrast ratios with horizontal lines indicating the maximum for the different configurations. The grey line denote the power law of exponent 1 in panels (b), (c), (e), and (f) and of exponent 2 in panel (g).

To estimate an upper limit of efficiency, we quantified the ideal contrast ratio that would be expected from the numerical generation of the hologram, which is simply the 2D Fourier transform of the phase mask. For a direct comparison, we took the patterns that were actually used in this experimental characterization. We found that the increase of background light and the corresponding decrease of contrast ratio when ROIs number increases are also expected by numerical results (Fig. 4f). Quantitatively speaking, the experimentally measured contrast ratios are higher than the one expected from holograms because of nonlinear 2P absorption that effectively boosts contrast.

One worry with the addition of the DMD might be related to the decrease in brightness. Pattern brightness when the DMD was used to reflect only the ROIs was reduced by about 25% (red and blue solid curves in Fig. 4g, respectively). But it should be noted that this decrease is small compared to the brightness that we would expect in a configuration without LCoS-SLM where a laser beam would be expanded onto the whole DMD. To mimic this situation, we generated a flat SLM pattern to illuminate the whole DMD chip uniformly (orange curve in Fig. 4e). For a given laser intensity and by extrapolating these curves, 2P absorption decreased by 5 orders of magnitude in this configuration. In other words, a laser intensity 200 fold higher was necessary to achieve the same level of fluorescence and even with a laser power of 34 W, we were not able to evoke sufficient 2P absorption for photoactivation. Also, this design generated patterns with reduced contrast ratios compared to holographic methods.

### Fast 2-photon stimulation

To assess the temporal characteristics of the optical stimulation, we used a rapid CMOS camera and imaged a fluorescent sample. In a first experiment, we reduced the field of view of the camera to several lines to achieve a frame rate of 10 kHz that matched the DMD framerate. We built a phase mask to generate 6 ROIs that were successively illuminated for 1 ms via updating the corresponding patterns on the DMD (Fig. 5a). This enabled us to verify that the temporal resolution of the set-up was only limited by the frame rate of the DMD and that no cross-illumination was generated by introducing the DMD between the LCoS-SLM and the objective.

**Fig. 5.**
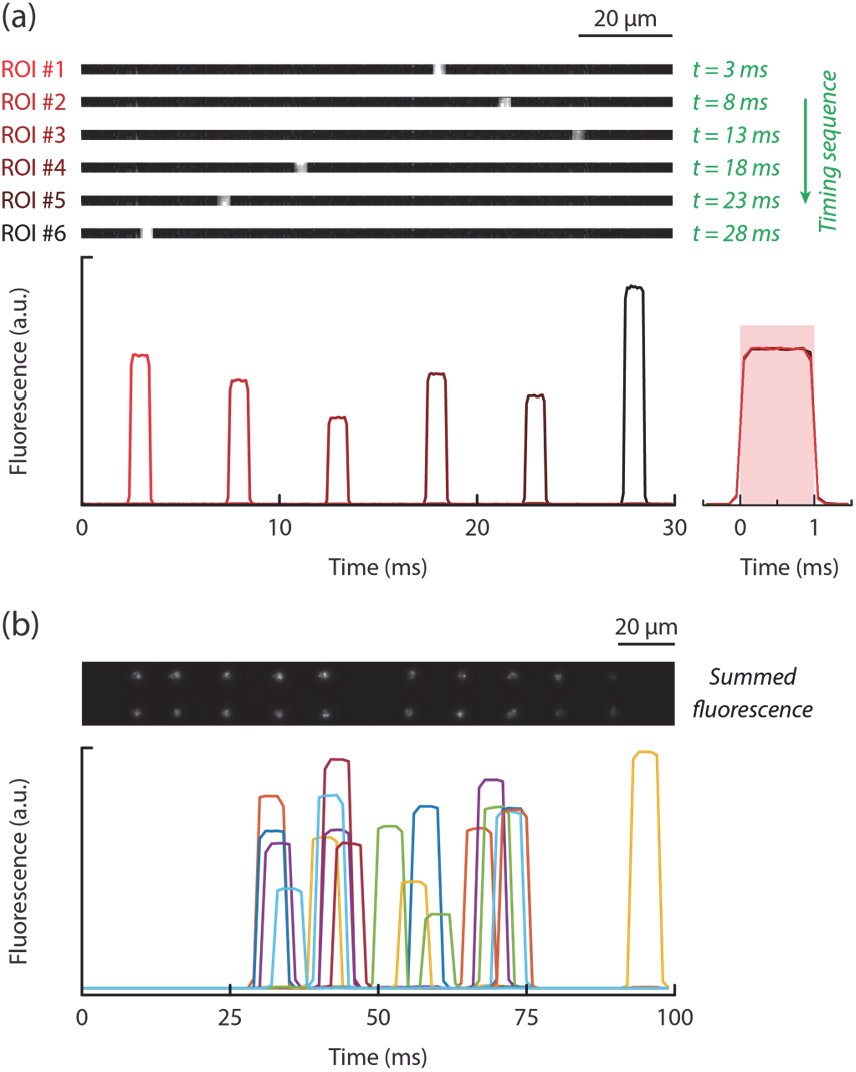
Temporal characterization of optical stimulation. (a) 6 ROIs illuminated consecutively for 1 ms by updating the DMD patterns (DMD and camera framerates: 10 kHz). Each image shows the fluorescence at a single point in time indicated in green on the right. Bottom graph shows the fluorescence signal of each ROI as a function of time, with a zoom around the period of stimulation on the right (traces were aligned to the stimulation time and normalized in terms of intensity). Note that the fluorescence rises from 5% to 95% in one acquisition frame (i.e. 0.1 ms). (b) 20 ROIs illuminated randomly for 5 ms by updating the DMD pattern (DMD framerate 10 kHz and camera framerate 1 kHz). The fluorescent image shows here the average light intensity during the duration of a whole sequence. Bottom graph shows the fluorescence signal of each ROI as a function of time.

In a second experiment, we increased the number of ROIs to 20 to make it more realistic regarding brain stimulation. The frame rate was decreased to 1 kHz to increase the field of view of the camera and the stimulation duration was set to 5 ms for each ROI (Fig. 5b). However, the DMD was still displaying patterns at 10 kHz and onsets had thus a 0.1 ms resolution. Also in this case, each ROI was faithfully activated without cross-illumination demonstrating that our approach permits to activate independently an arbitrary number of targets.

### Amplitude modulation

The DMD can be used as a temporal modulator but also as an amplitude modulator. To obtain the maximal intensity *I*_*max*_, all mirrors are turned ON within a single ROI. Because a typical ROIs (10 μm diameter disk at the sample plane) corresponds to ∼300 DMD pixels (one DMD pixel is 0.50 μm x0.53 μm at the sample plane), one can potentially select only a fraction *P* of the mirrors (Fig. 6a) to illuminate a given target with a fraction of the total intensity *P* · *I*_max_. To display spatially homogeneous patterns, we generated random patterns where each pixels within the ROI had a probability *P* to be ON. We updated these random patterns at the refresh rate of the DMD (i.e. 10 kHz). Therefore, the positions of ON pixels was changed at each frame to provide an average illumination intensity. This approach is of course applicable only if the biological process under study is slower than the DMD framerate (> 10 kHz), which is often the case.

**Fig. 6.**
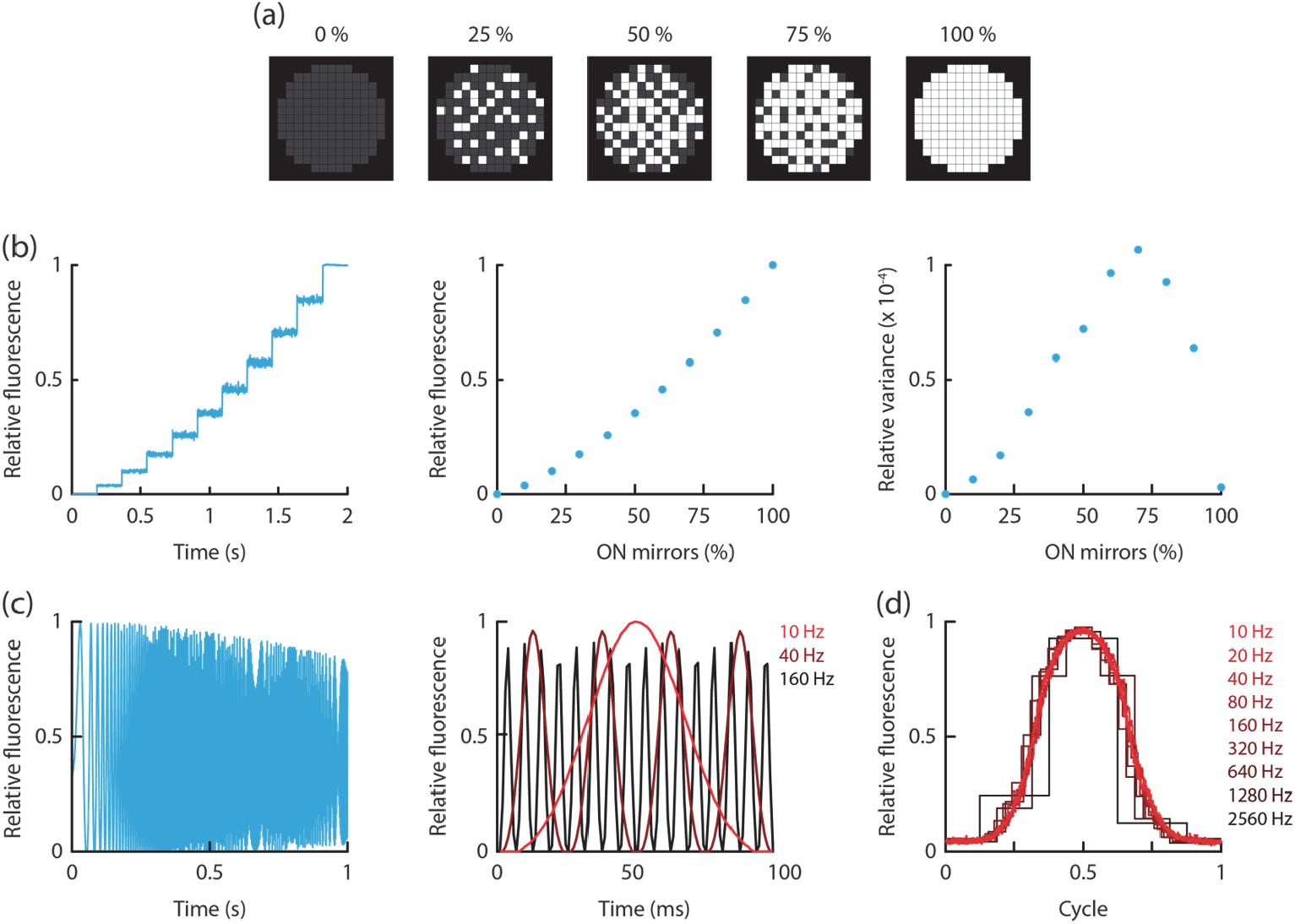
Using the DMD as a fast amplitude modulator. (a) Examples of DMD patterns with variable fractions of ON mirrors in order to modulate the average illumination amplitude of a single ROI. Note that in experiments, these patterns were randomly generated and varied at each DMD frame. (b) Step modulation. Relative fluorescence (left) of a 10 μm diameter ROI measured using the camera and plotted as a function of time when an increasing fraction of mirrors on the DMD were turned ON by steps of 10%. The average fluorescence (middle) and variance (right) were calculated during each step and plotted as a function of percentage of ON mirrors. (c) Frequency modulation measured with the camera. Chirp response for a modulating frequency varying between 1 and 500 Hz (left). Examples of sinusoidal modulation for 3 frequencies and plotted as a function of time. (d) Frequency modulation measured with the PMT and plotted as a function of cycle to display the frequency limit of the device. Note that all temporal traces displayed here were not averaged and correspond to single trials.

We verified the performance of the method by increasing the fraction of ON mirrors on a single ROI while measuring the integrated fluorescence on the ROI shape with the camera (Fig. 6b, left). We quantified the average fluorescence intensity for each fraction of ON mirrors (Fig. 6b, middle). The relation was monotonic but nonlinear because the size of a single DMD mirror on the sample plane (∼500 nm) is on the same order of magnitude than the microscope resolution. Thus, neighboring mirrors were not independent and nonlinear 2P effects were still present. However, a proper calibration permits to target the exact desired light intensity. We also observed fluctuations around the mean in the fluorescence trace (Fig. 6b, left), which were due to different realizations of the DMD patterns and not to measurement noise. For a given targeted intensity, fluctuations were low when mirrors are mostly ON or OFF and highest for intermediate value (Fig. 6b, right). Fluctuations could arise from two reasons: 1) variations in the number of ON pixels at each time step (because each pixels had a given probability to be ON), and 2) variations in the intensity of each pixel (because of speckles). The bell-shape variance makes sense because the intensity at each time step should follows a binomial distribution whose variance is proportional to *P*· (1 -*P*), where *P* is the probability of a mirror to be ON. Note that the peak was not observed for *P* = 0.5 again because of nonlinear 2P absorption. If needed, total noise could be reduced by fixing the percentage of ON mirrors at each frame instead of assigning an ON probability for each pixel but there would still be unavoidable fluctuations because of speckles.

We then turned to fast modulations of intensity. Using either a camera (Fig. 6c) or the fast PMT (Fig. 6d) to collect the average fluorescence intensity of the whole ROI, we established that a sinusoidal modulation of light intensity was possible up to the kHz frequency range. Taken together, this shows that this strategy enables to stimulate independently and with varying intensities an arbitrary number of ROIs with a frame rate only limited by the refresh rate of the DMD.

### Modulation of 3D patterns

In this section, we extend our stimulation method to patterns in 3 dimensions. First, we characterized the spatial footprint of patterns along the lateral and axial dimensions when the targets were at different axial positions (Fig. 7a). As expected, we observed an enlargement along the axial position when the ROI was away from the objective focal plane. Although, our approach is based on conjugating the amplitude pattern produced by the LCoS-SLM with the plane of the DMD, it is still possible to modulate a ROI even if it is not in the plane of the DMD as long as the DMD mask is sufficiently large and thus ROIs were not too close. To characterize the optimal mask on the DMD, we varied both the diameter of the disk on the DMD and the axial position of the ROI and measured the integrated fluorescence in the XY center plane (Fig. 7b). We observed that out-of-focus targets needed larger disks on the DMD to be efficiently relayed to the sample (Fig. 7c). As a result, if targets are within ± 100 μm in respect to the objective focal plane, mask diameters on the DMD do not need to be larger than ∼100 μm to obtain efficient light delivery and thus limited cross-activation. Interestingly, it is even possible to modulate two ROIs that are superimposed in XY but located at different axial planes if one target lies in the objective focal plane and the other is sufficiently separated.

**Fig. 7.**
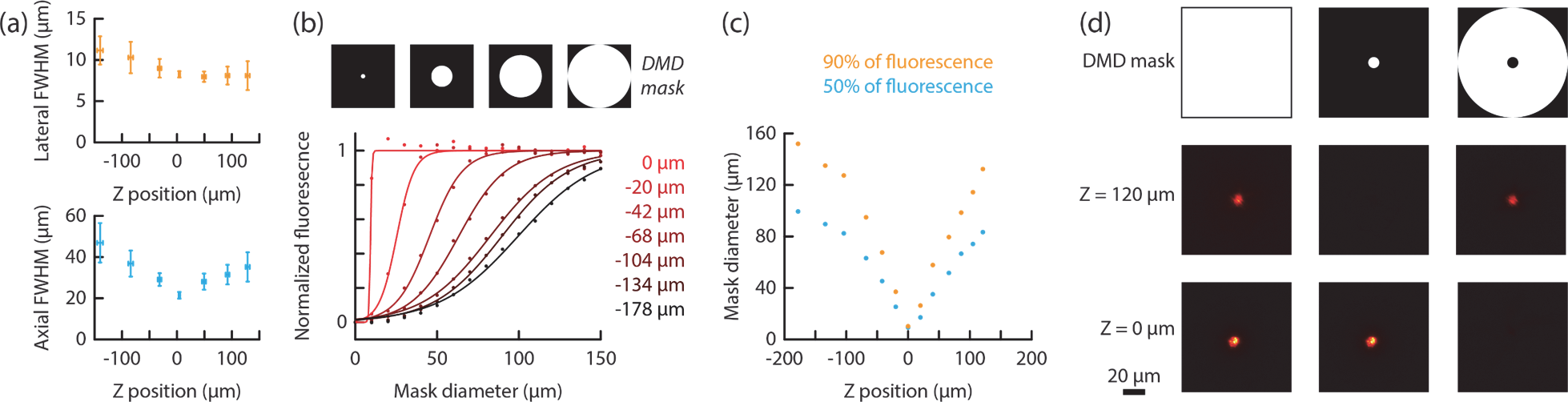
Modulation of patterns in 3 dimensions. (a) Spatial footprint measured as the full width at half maximum (FWHM) of the lateral (top) and axial (bottom) integrated intensities as a function of axial position of the target ROI (10 μm diameter, n = 16 ROIs on the same pattern). The DMD was fully reflective in this case. (b) Normalized fluorescent intensity as a function of DMD mask diameter for ROIs at different axial positions (from red to black). (c) Mask diameter on the DMD necessary to reach 50% (cyan) or 90% (orange) of the maximal fluorescence as a function of axial position of the target. (d) Intensity profiles in the XY plane at two different axial positions when the LCoS-SLM was generating 2 ROIs centered at these two axial locations. Three different DMD masks were used to select both ROI (left, fully reflective DMD), the ROI on the objective focal plane (center, small disk on the DMD) or the out of focus ROI (right, donut shape on the DMD).

In this condition, a disk on the DMD adjusted to the size of the ROI will only select the target in the focal plane (Fig. 7d, middle). Conversely, a donut shape on the DMD will only opt for the out-of-focus target (Fig. 7d, right).

We further characterize the 3D capability of our system by constructing more complex multidimensional patterns. To target 4 different planes, we divided the grid of the LCoS-SLM in 4 sub-grids that were each targeting a different plane (Z = -95, -40, 45, 100 μm). Pixels from each sub-array were uniformly distributed to maximize homogeneity of the light source intensity. We designed disks on the DMD that had a diameter of about 60 μm compared to the 8 μm of the actual ROI. Compared to a case where all the mirrors of the DMD are ON (Fig. 8a), the pattern quality was not significantly affected when only mirrors corresponding to the different ROIs where ON (Fig. 8b). Thus, we could selectively erase 1 or more ROIs (Fig. 8c-d) even if the image generated by the LCoS-SLM is out of the DMD focus. Importantly, the selective modulation of the patterns in 3D could be done at the DMD framerate.

**Fig. 8.**
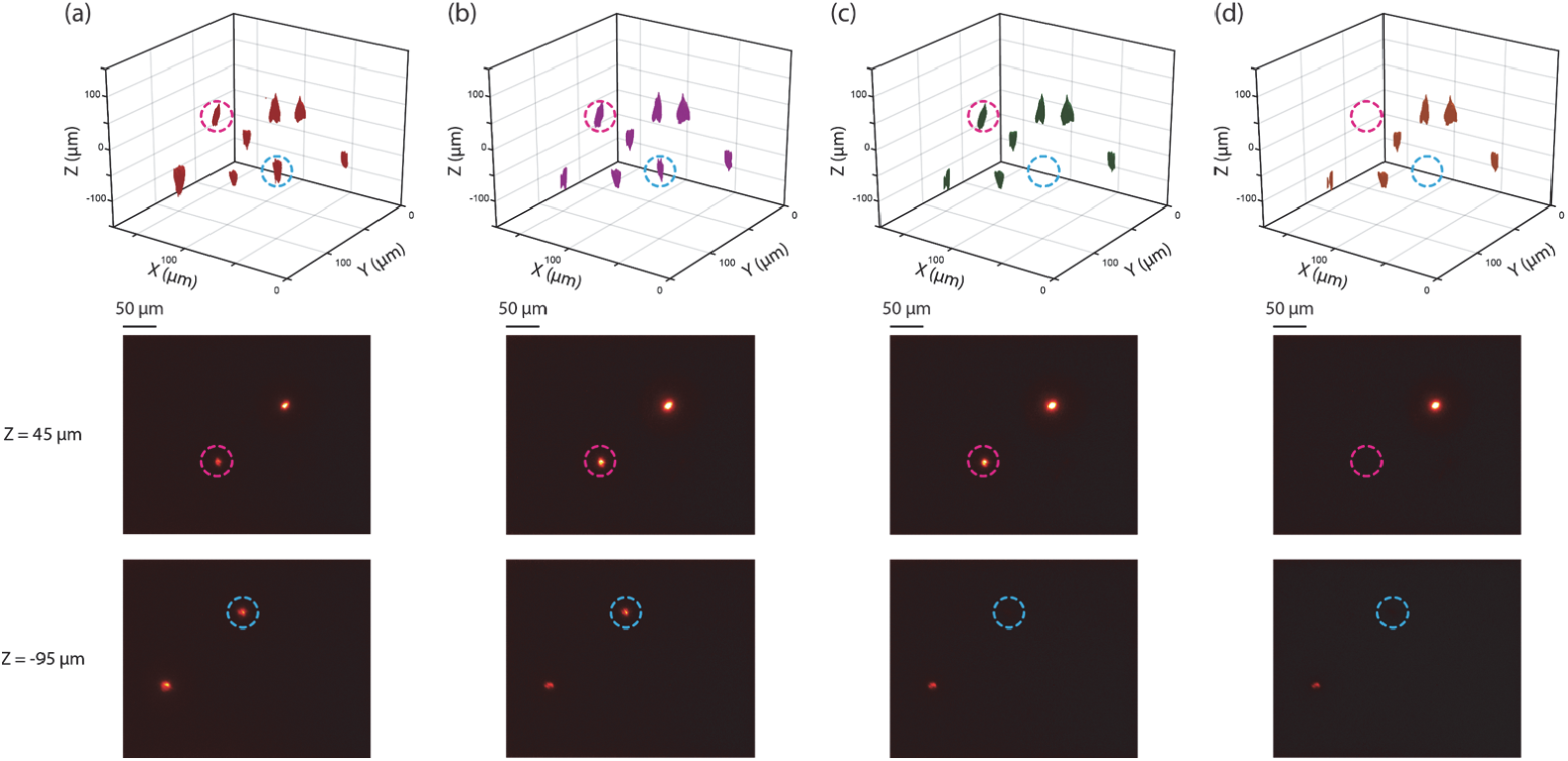
Selective modulation of patterns in 3 dimensions. (a) 3-dimensional reconstruction (top) of the optical patterns when all the mirrors of the DMD where in the ON position. Two selected planes are shown (bottom). (b-d) Same as in (a) when only the mirrors corresponding to the 8 ROIs where ON (b) or when only 7 (c) or 6 (d) ROIs were ON on the DMD.

### Physiological assessment of spatial characteristics

To characterize the spatial performance of the optical stimulation functionally, we grew primary cortical neurons onto a glass coverslip and expressed the light-gated ion channel channelrhodpsin ChR2(H134R) using viral transfection (see Methods). We monitored membrane potential using patch-clamp recordings (Fig. 9a). Cells had intrinsic properties similar to published data (Table 1) (Barral and Reyes 2016). We stimulated neurons either using 2P light patterning as described above or with the fiber-coupled LED of the 1P fluorescent light path where an iris was placed on the conjugated plane of the sample (Fig. 1). Light intensity for 1P activation (5-10 mW·mm^-2^) was close to the saturation level and was distributed over a disk of 60 μm diameter. For 2P activation, non-saturating power of 60-120 W·mm^-2^ (i.e. 10-20 mW in a 15 μm disk or 60-120 μW·μm^-2^) was used to avoid heating cells substantially. Upon light activation, light-sensitive channels on the neuron’s membrane open, leading to a photocurrent and thus neuron’s depolarization from its resting membrane potential (Fig. 9b). Neurons were significantly more depolarized using 1P than with 2P stimulation (Table 1) mostly because of the larger illumination spot for 1P than for 2P stimulation. Single photon activation is also often more efficient and has been observed for a variety of opsin variants even at saturating light amplitude (Rickgauer and Tank 2009, Andrasfalvy, Zemelman et al. 2010, Prakash, Yizhar et al. 2012). However, recent advances in biochemical engineering have now permitted to produce much more efficient opsins for 2P activation both in terms of kinetics and photocurrent (Sridharan, Gajowa et al. 2022).

**Table 2.**
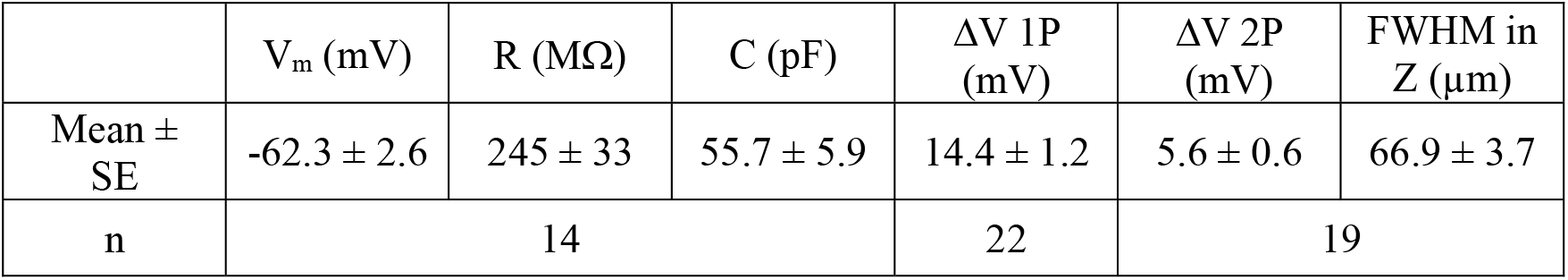
Electrophysiological characterization of neuron intrinsic properties and light stimulation

| | $V_m$ (mV) | $R$ ($M\Omega$) | $C$ (pF) | $\Delta V$ 1P (mV) | $\Delta V$ 2P (mV) | FWHM in Z ($\mu\text{m}$) |
| --- | --- | --- | --- | --- | --- | --- |
| Mean ± SE | $-62.3 \pm 2.6$ | $245 \pm 33$ | $55.7 \pm 5.9$ | $14.4 \pm 1.2$ | $5.6 \pm 0.6$ | $66.9 \pm 3.7$ |
| n | 14 |  |  | 22 | 19 |  |

**Fig. 9.**
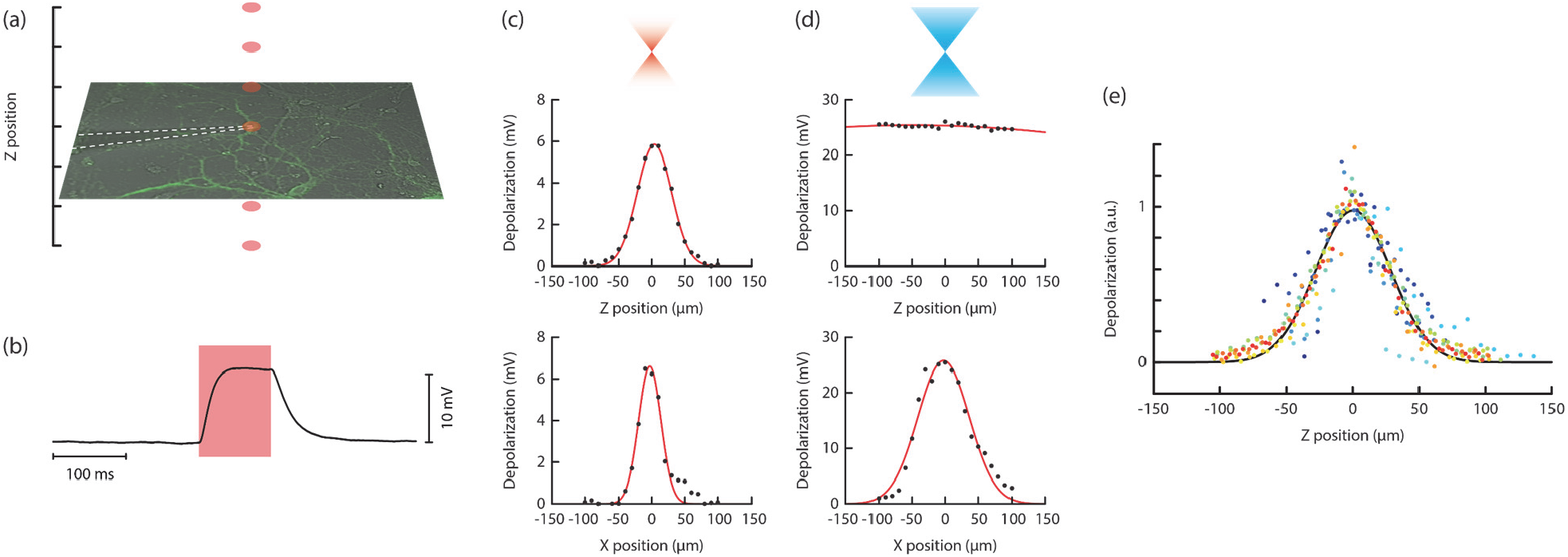
Functional characterization of the spatial precision of holographic stimulation. (a) Experimental design where patch-clamp recording was made from a cortical neuron in culture. (b) Example data of a depolarizing step upon 2P activation. (c-d) Steady voltage depolarization as a function of the axial (top) and lateral (bottom) position of the stimulation spot for 2P (c) and 1P (d) stimulations. Except for 1P axial sensitivity, experimental data points were fitted by a Gaussian curve. (e) Population data showing the normalized voltage depolarization as a function of the axial position of the stimulation spot for 2P activation (n = 19 neurons). For each individual neuron, fit parameters were used to normalize the maximal depolarization to 1 and center the Gaussian curve to Z = 0. An axial precision of 67 μm (FWHM) for a spot of 15 μm diameter was found.

Here, we used this approach to describe only the spatial resolution of the setup because the slow kinetics of the opsin did not permit to fully appreciate the fast temporal performances of the microscope. Further experiments using fast opsins would be valuable to assess the temporal characteristics of the setup. We measured the depolarization of single neurons as a function of the position of the stimulation pattern. Depolarization dropped by 50% when the stimulation region was moved within the sample plane by about 54 μm and 32 μm for 1P and 2P stimulation, respectively (Fig. 9c-d). Note that the lower resolution for 1P stimulation is mainly due to the iris size that had a diameter of about 60 μm on the sample plane, compared to the 15 μm spot diameter that were generated with the LCoS-SLM. Upon 2P stimulation, depolarization was decreased by 50% when the focus was moved away by 34 μm (Table 1, Fig. 9c-e), a value comparable to published data using conventional holography (Pegard, Mardinly et al. 2017, Shemesh, Tanese et al. 2017) given that our stimulation spots were 15 μm and had thus broader axial tuning (axial FWHM is 25 μm for 15 μm spots; Fig. 3e). Notably, these profiles could be substantially sharpened along the optical axis using temporal focusing (Andrasfalvy, Zemelman et al. 2010, Papagiakoumou, Anselmi et al. 2010, Pegard, Mardinly et al. 2017).

## Discussion

### Relation to previous work and potential improvements

A DMD has already been used for a rapid switching of targets and has found applications in population calcium imaging using multiplexing (Ducros, Goulam Houssen et al. 2013) or in multi-site photolysis of caged neurotransmitters (Go, To et al. 2013). Here, we advance this approach by decreasing the background illumination, demonstrating the ability to modulate the amplitude of each target independently, and assessing the 3D capability of the system. Further improvements on the axial resolution would be effective with the use of temporal focusing. For this, a diffraction grating needs to be introduced in conjugated plane of the DMD. It could be either a reflective gratings if the optical path permits or a volume phase holographic transmission grating. Yet, one have to keep in mind that the DMD is also dispersive because it is an array of tilted mirrors analog to a blazed grating. As such, DMD has been used as the dispersive element in a temporal focusing setup (Yih, Hu et al. 2014) but could nonetheless be coupled with a diffraction grating for 2 photon application (Li, Cheng et al. 2012).

An alternative approach to increase the temporal resolution of a LCoS-SLM based system is multiplexing. This has been done by incorporating two SLMs along the same photostimulation path, enabling temporally multiplexed ensemble stimulation (Marshel, Kim et al. 2019). Each SLM was used alternatively by switching the polarization of the incident beam. Thus, this approach can only increase the speed limit by a factor of 2 for a single wavelength system. An elegant study showed that multiplexing is also possible using a single LCoS-SLM. By sequentially illuminating discrete bands on the LCoS-SLM using galvo-based scanning, kHz rates were achieved (Faini, Tanese et al. 2023). However, both approaches necessitate more complex hardware and software implementations and are not compatible with fast real-time control because of the computation time necessary to design the collection of phase masks. Depending on the application and on the tradeoff between temporal resolution, contrast ratio, 3D capability, real-time update of patterns, and simplicity, the end-user has a variety of options available.

### Potential benefits of a DMD-based system

All-optical methods, where presynaptic neurons are optogenetically activated and postsynaptic neurons activity is read by calcium of voltage fluorescent indicators, allow to probe synaptic connections efficiently in the intact brain (Packer, Russell et al. 2015, Baker, Elyada et al. 2016, McRaven, Tanese et al. 2020). Development of compressed sensing algorithms permits to explore several presynaptic neurons at the same time and to increase the rate at which the existence and strength of synapses are screened (Triplett, Gajowa et al. 2022). However, to search for relevant connections, this class of algorithms needs to update the stimulation patterns in real time and the generation of phase mask appeared to be the limiting step. By selectively activating a subset of the potential targets and by modulating the amplitude of these ROIs to titrate the resulting activation, our optical design integrating a DMD could be used to probe synaptic connections with graded stimulation intensities without the need of computing a new phase mask for each stimulation pattern.

The increase of contrast ratio that we report here would be valuable for such experiments were cross-activation can produce confounding artifacts. This improvement of contrast is due to the imperfect spatial pattern generated by the LCoS-SLM where light is also diffracted to unwanted regions. By discarding part of this diffracted light, the DMD permitted to reduce background illumination. With our settings, 128 ROIs could be displayed simultaneously while keeping a contrast ratio above 97% as measured in the XY plane.

### Need for temporal resolution

A temporal resolution of 0.1 ms might not seem necessary at first sight because neuronal spike latency upon light stimulation is usually around 3-6 ms (Yizhar, Fenno et al. 2011, Ronzitti, Conti et al. 2017, Mardinly, Oldenburg et al. 2018, Sridharan, Gajowa et al. 2022). However, spike timing precision is well below latency values. Using the high-speed variant opsin Chronos, fast spiking interneurons could trigger spikes with a temporal precision of 0.2 ms upon 2P activation (Ronzitti, Conti et al. 2017). Sub-millisecond time jitter was also achieved with the newly developed opsin family ChroME (Mardinly, Oldenburg et al. 2018, Sridharan, Gajowa et al. 2022). ChroME variants have activation spectra centered around 1000 nm, a slightly higher wavelength than Chronos, which make them ideal for stimulation using Yb:YAG fiber lasers that can provide the necessary power to target a large number of cells in parallel and study how spatiotemporal patterns of neuronal activation are both necessary and sufficient for encoding information in the brain.

Spike timing with millisecond precision is important in many physiological processes such as spike-timing dependent plasticity (Bi and Poo 1998), song generation in the bird song (Hahnloser, Kozhevnikov et al. 2002), sensory processing (Rose, Brugge et al. 1967), or hippocampal memory replay (Carr, Jadhav et al. 2011). With the aim at reproducing precise neuronal sequences, our approach coupled to newly developed opsins provides a valuable resource to tackle fundamental questions about the role of temporal information in neuronal coding.

### Extension to 3D patterns

The ability to produce patterns in 3 dimensions is important for applications in biological tissues. An axial resolution of the 2P stimulation of the order of 67 μm for a spot of 15 μm diameter was found. This value could be reduced to ∼10 μm with the addition of temporal focusing technique described in several studies (Andrasfalvy, Zemelman et al. 2010, Papagiakoumou, Anselmi et al. 2010, Pegard, Mardinly et al. 2017). It is thus expected that the single cell resolution can be achieved in complex biological tissues. This resolution is often necessary when dealing with specimens in 3 dimensions to avoid unwanted cross-activations.

Previous approaches used either holography (Pegard, Mardinly et al. 2017), tunable lenses (Yang, Carrillo-Reid et al. 2018) or AOD (Ricci, Marchetti et al. 2022) to target different planes, all of which did not provide a sufficient temporal resolution to activate a large population of neurons in 3D simultaneously or with precise temporal patterns. Our optical design alleviates this limitation by using the DMD as a temporal modulator of each ROI independently. For 3D applications, we showed that temporal patterns could be modulated by the DMD as long as the different patterns are not too close 1) in XY if they are close in Z, or 2) in Z if they are close in XY (Fig. 7). Here, we were able to control specifically individual ROIs from different planes that were separated by up to 200 μm which represents a volume relevant for the study of cortical processes for example.

## Acknowledgments

We thank Nicolas Michalski and Boris Gourévitch for lending us the CMOS camera and Brice Bathellier for the virus sample. This work was supported by a Human Frontier Science Program Career Development Award (CDA00009/2017-3), by the CNRS Momentum program and by the Fondation pour l’Audition (FPA IDA01). We acknowledge the support of the Fondation pour l’Audition to the Institut de l’Audition.

## Disclosures

The authors declare that their principal affiliations are with not-for-profit organizations and that they have no potential conflict of interest in the publication of this article.

## Data availability

The data that support the findings of this study are not publicly available at this time but may be obtained from the corresponding authors upon reasonable request. Data acquisition (Labview) and analysis (Labview or Matlab) software used in this paper are described in the Methods and will be available upon request.

## Notes

### Competing Interest Statement

The authors have declared no competing interest.

### Summary of Updates

This version of the manuscript contains more details about the experimental design and further characterization of the spatiotemporal performances.

